# First detection of *Anopheles stephensi* in Ghana using molecular surveillance

**DOI:** 10.1101/2023.12.01.569589

**Authors:** Yaw A. Afrane, Anisa Abdulai, Abdul R. Mohammed, Yaw Akuamoah-Boateng, Christopher Mfum Owusu-Asenso, Isaac K. Sraku, Stephina A. Yanney, Keziah Malm, Neil F. Lobo

**Author notes:** **Address for Correspondence** Yaw A. Afrane, (YAA).

## Abstract

The invasive *Anopheles stephensi* mosquito has been rapidly expanding in range in Africa over the last decade, spreading from the Indian sub-continent to several East African countries (Djibouti, Ethiopia, Sudan, Somalia and Kenya) and now in West Africa, Nigeria. The rapid expansion of this invasive vector poses a major threat to current malaria control and elimination efforts. In line with the WHO’s strategy to stop the spread of this invasive species by enhancing surveillance and control measures in Africa, we incorporated morphological and molecular surveillance of *An. stephensi* into routine entomological surveillance of malaria vectors in the city of Accra, Ghana. Here, we report on the first detection of *An. stephensi* in Ghana. *An. stephensi* mosquitoes were confirmed using PCR and sequencing of the ITS2 regions. These findings highlight the urgent need for increased surveillance and response strategies to mitigate the spread of *An. stephensi* in Ghana.

## Background

*Anopheles stephensi* is an invasive mosquito species originating from parts of Southeast Asia and the Arabian Peninsula (1). The ability of this species to utilize artificial containers for larval sites has made this vector capable of thriving in urban areas, setting them apart from other major malaria vectors that primarily breed in rural areas (2). *An. stephensi* is capable of transmitting both *P. falciparum* and *P. vivax* (1,3). Over the last decade, *An. stephensi* has been expanding in range and has now been documented in several countries in Africa (4). It was first detected in Djibouti, the Horn of Africa in 2012, where it was implicated in an urban malaria outbreak (5). It was also detected in Ethiopia in 2016 and 2018, where it is well-established in eastern Ethiopia (6,7). *An. stephensi* was subsequently detected in Sudan (2016), Somalia (2019), Nigeria (2020) and Kenya (2023) (4,5,7–9).

The rapid expansion of *An. stephensi* in sub-Saharan Africa (SSA) which has the highest burden of malaria globally, is a major public health concern. The spread of this invasive species could lead to high malaria transmission in urban areas though malaria is typically a rural disease. In Djibouti, *An. stephensi* mosquitoes are thought to be responsible for an increase in malaria incidence, from 1 to 4 cases in 2013 to 49.8 cases/1,000 persons in 2019 (10). With over 40% of the population in SSA living in urban areas, the spread of *An. stephensi* into these receptive areas will currently put about 126 million people at risk of malaria (2,4). Also, this invasive vector has been found to be resistant to insecticides further increasing the risk of malaria transmission in combination with limiting intervention efficacy (11–13). *An. stephensi* mosquitoes from Somalia were found to be resistant to several insecticide classes, especially pyrethroids (13).

The World Health Organization issued an initiative in 2022 aimed at strengthening surveillance, increasing collaborations and prioritizing research to help stop the spread of *An. stephensi* in SSA and find strategies to combat or eliminate the vector in areas that have been invaded (4). Morphological and molecular surveillance of *An. stephensi* were incorporated into routine entomological surveillance of malaria vectors in the city of Accra, Ghana, following the WHO initiative, that seeks to take coordinated action to limit the spread of this invasive species by improving surveillance and control efforts in Africa (4). This study outlines the entomological surveillance that documents the identification of this invasive species in Ghana.

## Methods

### Study Sites

Sampling was conducted in 8 sites within the city of Accra, Ghana, as part of routine entomological surveillance from January 2022 to July 2022. These sites were categorized to represent different environments and socio-economic status; irrigated urban farming (IUF) sites (Tuba and Dzorwulu), lower socioeconomic (LS) sites (Nima and Chorkor), middle socioeconomic (MS) sites (Dansoman and Teshie) and high socioeconomic (HS) sites (East Legon and Cantonment). Tuba (5° 30’ 47”N 0° 23’ 16” W) and Dzorwulu (5°36′53″N 0°12′03″W) are sites where irrigated farming is practised all year round leading to the creation of mosquito breeding sites. Socio-economic sites were classified based on their population, housing structures and the availability of proper drainage and sanitation systems. Low socioeconomic sites, Nima (5° 35′ 0″ N, 0° 12′ 0″ W) and Chorkor (5°31′39″N 0°13′55″W) are densely populated slums with poor sanitation and inadequate drainage systems. Dansoman (5° 33′ 0″ N, 0° 16′ 0″ W) and Teshie (5° 35′ 0″ N, 0° 6′ 0″ W) are middle socioeconomic sites with more standard residential structures with well-designed drainage and sanitation systems but poorly managed. High socioeconomic sites, Cantonment (5° 35′ 10″ N, 0° 10′ 35″ W) and East Legon (5°38’16.39”N, 0°9’40.33”W) have proper housing structures with good sanitation and drainage systems. Accra is the capital city of Ghana and it is the most populous. Accra lies in the coastal savannah zone of Ghana, with an annual mean temperature of 26.5 °C and an average annual precipitation of 787 mm. Figure 1 shows a map of the routine surveillance sites.

**Figure 1:**
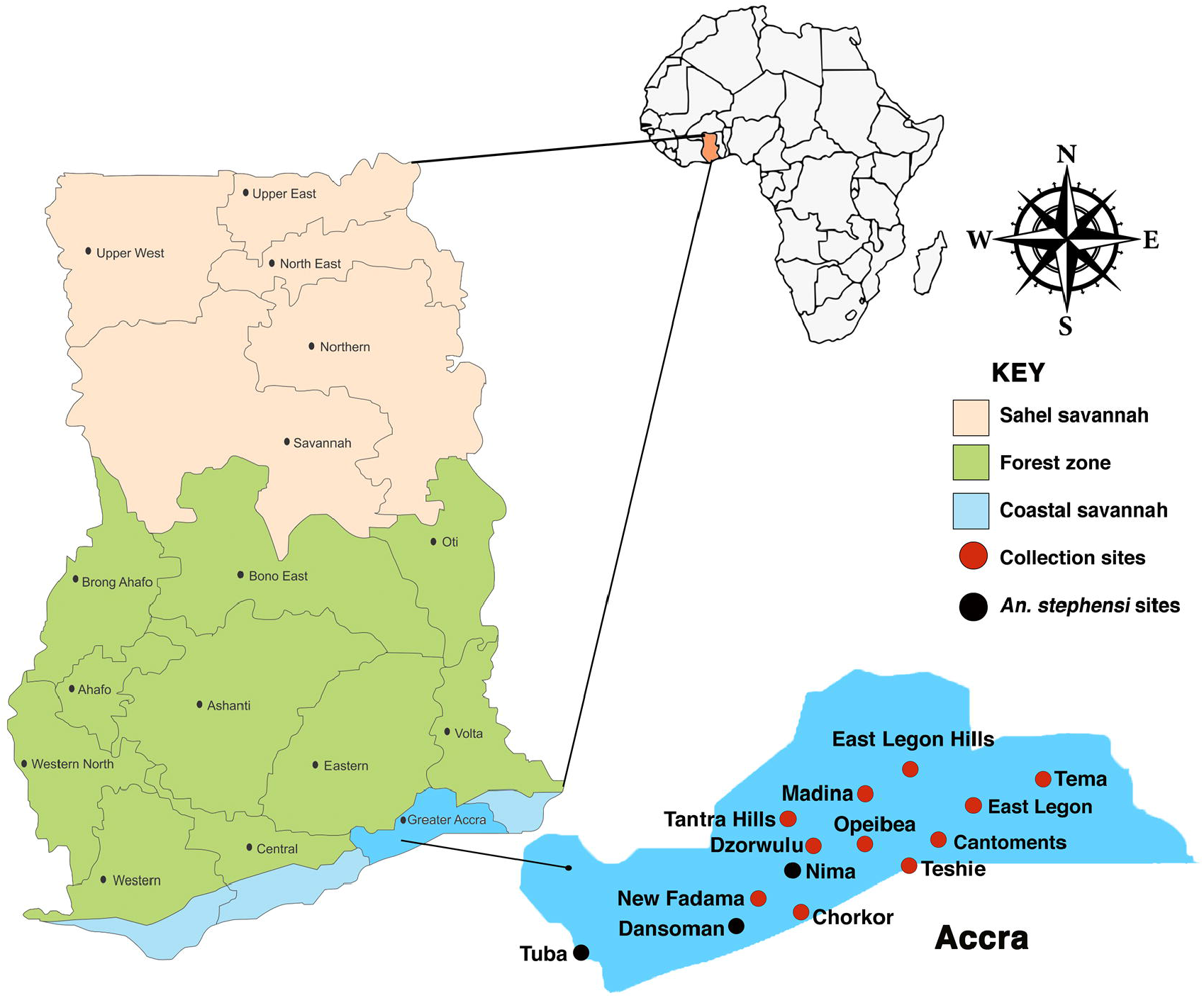
Routine entomological surveillance sites in Accra, Ghana

### Larval Habitat Characterization

Larval habitats identified in each site were grouped into two; natural habitats and man-made habitats. The man-made habitats included ditches, footprints, tyres and tyre tracks while natural habitats included swamps, furrows and natural ponds. The land-use type where the larval habitats were found was recorded. The geographical coordinates of each larval habitat were recorded using a GPS device (Garmin eTrex 10 Worldwide Handheld GPS Navigator).

### Larval mosquito sampling and densities

Larval sampling was conducted for all potential breeding sites using the standard WHO dippers and small ladles for smaller habitats (14). The total number of dips was recorded as described by Hinne *et al*. (14). The number of larvae and pupae was recorded, and the larval density was calculated as the ratio of the number of larvae collected per dip (14,15). Larval sampling was done in every site monthly for the dry (February – March) and rainy (June – July) seasons of 2022. Larval samples were transported to the insectary at the Department of Medical Microbiology, University of Ghana Medical School, where they were raised into adults for morphological identification.

### Morphological and molecular identification of mosquito samples

Adults raised from larvae sampled were morphologically identified to species using their palps, wings, abdomen and legs using the keys of Nagpal and Sharma (16) and Coetzee (17). DNA was extracted from the mosquito legs using the alcohol precipitation method (18). PCR amplifications were carried out to detect *An. stephensi* using primers targeting the ITS region based on previously described protocols by Singh et al. (19). Members of the *An. gambiae s*.*l* complex were further identified by PCR using the extracted DNA as the template. Four sets of oligonucleotide primers (*An. gambiae, An. arabiensis, An. melas* and universal primer) were used in the PCR for the identification of members of the *Anopheles gambiae* s.l species complex (20). *Anopheles gambiae* s.s *and An. coluzzii* were distinguished by PCR-RFLP using previously described protocols (21).

### Molecular Species Identification - Sequencing

After PCR, mosquitoes that did not produce bands indicative of the *An. gambiae* complex (n=11) were subjected to Sanger sequencing of the ITS2 regions and analysed based on comparisons to the NCBI database (22).

## Results

### *Anopheles* larval densities in different habitat types across different sites

Ten (10) different habitat types were encountered during the larval sampling. The highest larval density during the dry and wet seasons was observed in drainage ditches from Chorkor (9.72 larvae/dip) and swamps in Teshie (20.3 larvae/dip) respectively. Drainage ditches were consistently productive across almost all the sites in both seasons. The most productive habitat type across all the sites was drainage ditches. However, habitat types such as footprints, swamps and tyre tracks also recorded low to high larval densities in some of the sites (0.25 to 20.3 larvae/dip). In Tuba, Nima and Dansoman, where *An. stephensi* mosquitoes were found, and some of the more productive habitats were drainage ditches (1.45 to 8.39 larvae/dip) and tyre tracks (0.77 to 14.96 larvae/dip) (Table 1). Figure 2 shows habitats where *An. stephensi* mosquitoes were found. *An. gambiae s*.*l*. larval density was significantly associated with season (*t* = 4.14, *P* = 0.00).

**Table 1:**
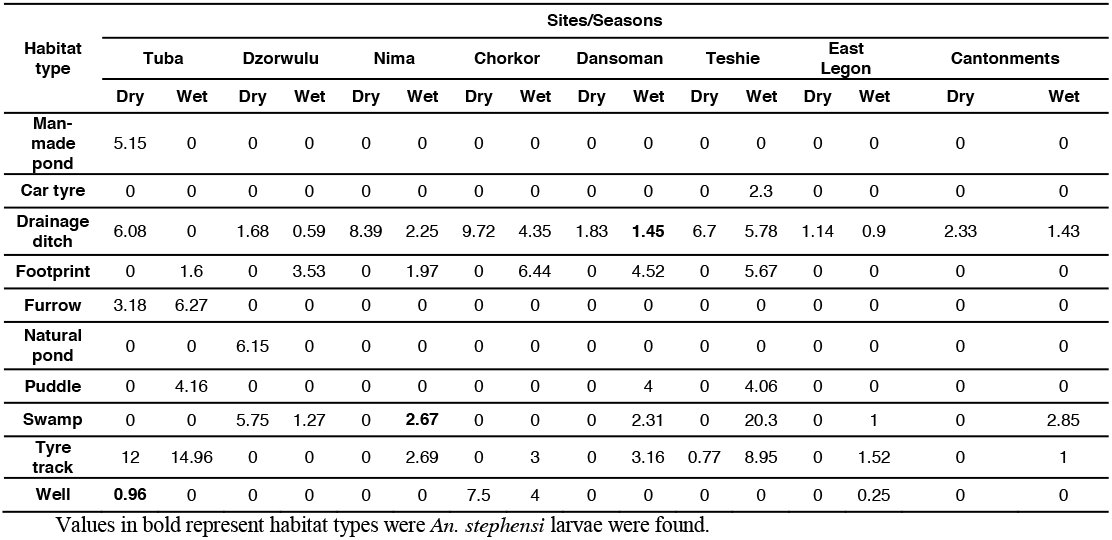
*Anopheles* larval density in the dry and rainy seasons.

**Figure 2:**
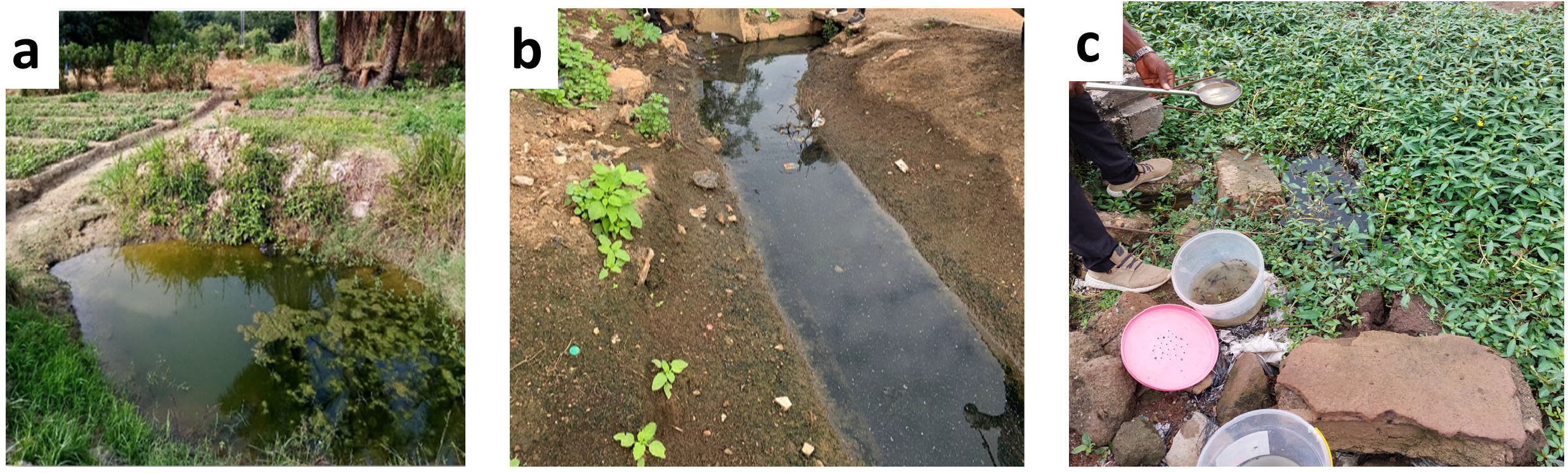
Habitats were *An. stephensi* larvae were found. **a** Dug-out well (Tuba), **b** drainage ditches (Dansoman), **c** swamp (Nima)

### Species distribution of *Anopheles* mosquitoes

A total of 1169 mosquitoes obtained from the larval sampling were identified using morphological keys and PCR methods for speciation. Out of this number, 551(47.13 %) were *An. gambiae s*.*s*, 582 (49.79 %) *An. coluzzii* and 32 (2.74%) Hybrids. Four samples (0.34 %) were identified as *An. stephensi* using a modified PCR-based method by Singh et al. (19) and sequencing (22)(Table 2). Results from the NCBI blast showed that the *An. stephensi* samples had 100% sequence similarity with *An. stephensi* voucher A268 5.8S ribosomal RNA gene and internal transcribed spacer 2 (GenBank: MH650999.1) (Table 3).

**Table 2:** *Anopheles* larvae species distribution across different sites.

| Site | Site Category | Species, no. (%) |  |  |  |  |
| --- | --- | --- | --- | --- | --- | --- |
|  |  | *An. gambiae* | *An. coluzzii* | Hybrids | *An. stephensi* | Total |
| **Tuba** | **IUF** | 197 (61) | 116 (35.9) | 8 (2.5) | 2 (0.6) | 323 (100) |
| **Dzorwulu** |  | 5 (31.3) | 11 (68.7) | 0 | 0 | 16 (100) |
| **Nima** | **LS** | 67 (33.5) | 120 (60) | 12 (6) | 1 (0.5) | 200 (100) |
| **Chorkor** |  | 17 (29.3) | 41 (70.7) | 0 | 0 | 58 (100) |
| **Dansoman** | **MS** | 7 (7.1) | 84 (85.7) | 6 (6.1) | 1(1.1) | 98 (100) |
| **Teshie** |  | 166 (46.62) | 186 (52.2) | 3 (1.2) | 0 | 355 (100) |
| **East Legon** | **HS** | 77 (77.7) | 19 (19.3) | 3 (3) | 0 | 99 (100) |
| **Cantonment** |  | 15 (75) | 5 (25) | 0 | 0 | 20 (100) |
| **Total** |  | **551 (47.13)** | **582 (49.79)** | **32 (2.74)** | **4 (0.34)** | **1169 (100)** |

**Table 3:** Sequencing results of suspected *An. stephensi* samples.

| Sample ID | ITS2 Contig | NCBI blast result | GenBank accession number of best match | %Identity match | Final Species ID | GenBank accession numbers |
| --- | --- | --- | --- | --- | --- | --- |
| DN 035 | 283 | *An. stephensi* voucher | MH650999.1 | 100% | *An. stephensi* | OR711900 |
| TP 002S | 283 | *An. stephensi* voucher | MH650999.1 | 100% | *An. stephensi* | OR711899 |

## Discussion

The invasion of *An. stephensi* in sub-Saharan Africa, which bears the world’s highest malaria burden, represents a significant concern for public health. This is because of their ability to thrive in urban areas and transmit both *P. falciparum* and *P. vivax*. Here we report the first detection of *An. stephensi* in Ghana using molecular surveillance. *An* .*stephensi* was found in larval mosquito samples from urban areas of Accra, Ghana, specifically Tuba, Dansoman and Nima.

While the vector’s spread could have occurred through land borders, air travel, or seaports, it is noteworthy that in Ghana, it was discovered at considerable distances from these points of entry, suggesting possible introduction via long-distance migration (Atieli et al 2023), local transportation, and/or human activities. Similar studies in Eastern Ethiopia have reported the collection of *An. stephensi* samples far inland along transportation routes that are not proximate to any seaport entry, underscoring the role of long-distance migration, local transportation, and human activities in driving the dispersal of this invasive species (23). This highlights the need to expand surveillance efforts to determine the distribution and spread of *An. stephensi* in Ghana. It is likely that this invasive species may have spread to other parts of Accra as well as other regions of Ghana.

*Anopheles stephensi* is known to breed in various types of larval habitats, including manmade water containers such as plastic tanks, cisterns, barrels, discarded tires, and plastic receptacles, as well as freshwater pools such as stream margins and irrigation ditches. Remarkably, in this study, *An. stephensi* was found breeding in habitats distinct from the typical ones observed in Asia and East Africa (10,24). In Ghana, this vector was identified in dug-out wells within irrigated vegetable farms and roadside ditches. Additionally, it was observed to breed alongside *An. gambiae s*.*s* and *An. coluzzii*, whereas it is commonly associated with Aedes mosquitoes.

Expanding surveillance efforts for *An. stephensi* in both urban and rural areas should be a primary focus of Ghana’s National Malaria Elimination Program. Such efforts are crucial to curbing the dissemination of this invasive species within Ghana, which could potentially elevate malaria prevalence in Accra, traditionally considered a low malaria transmission zone within Ghana(25). The rapid expansion of *An. stephensi* also raises the risk of its colonization in rural regions of Ghana, where malaria prevalence is already high, resulting in intensified malaria transmission, disease morbidity, and mortality. Incorporating molecular-based detection tools into surveillance systems is paramount for the early detection of invasive malaria vectors, preventing their adaptation and local establishment(8).

## Conclusion

The first report of the invasion of *An. stephensi* in Accra, Ghana, represents a major public health concern, given the heightened risk of urban malaria outbreaks. It is imperative to reinforce surveillance and response strategies in both rural and urban settings across Ghana, with specific attention directed towards *Anopheles stephensi*, to mitigate the spread of this invasive species.

## Acknowledgement

This study was supported by grants from the National Institute of Health (NIH: R01 A1123074 and D43 TW 011513).

YAA, KM and NFL were responsible for the study design, supervised the data collection and contributed to the writing of the manuscript. AA, ARM, YAB, CMO-A, SAY and IS performed the data collection, laboratory work and analysis. AA, YAA and NFL drafted the manuscript. All the authors read and approved the final manuscript.

## Biography

Yaw A. Afrane is a Professor of Vector Biology at University of Ghana, Accra, Ghana. His research focus on vector and parasite biology and epidemiology with over 15 years of research experience in vector-borne diseases.

